# DigitalPedon: A Novel Digital Twin Framework for Soil Profile Monitoring and Global Soil Data Interoperability

**DOI:** 10.64898/2026.05.05.722891

**Authors:** Ali Youssef, Nasem Badreldin

## Abstract

The Digital Pedon (DP) is an open-source Python framework that represents a soil profile as a continuously updated digital twin, bridging three persistent gaps in soil science: disconnected models and observations, cross-database interoperability, and the inference gap between raw sensor signals and agronomically meaningful variables. Integrating real-time sensor streams, model-based solver chains (Model-Zoo), GLOSIS-compliant ontology mapping, and a novel LLM agentic interface layer enabling natural language soil queries, the DP supports applications spanning precision agriculture, digital soil mapping, and environmental sustainability assessment. Four proof-of-concept experiments confirm automatic profile initialisation fidelity, solver chain consistency, ontology compliance, and user-defined solver extensibility.

## 1. Motivation and significance

Global soil health is one of the most important assets to a sustainable food production and climate resilience [1], [2]. Soils are dynamic systems whose physical, chemical, and biological properties continuously vary in response to environmental drivers (e.g., precipitation, drought), land management (e.g., tillage, irrigation), and ecological interactions (e.g., microbial decomposition, mycorrhizal symbiosis). Yet most available soil information systems still represent soils as static entities, relying on infrequent field surveys, laboratory analyses, and profile descriptions that capture conditions at a single point in time using invasive methods. While these measurements provide essential information on soil properties and classification, they cannot capture the rapid shifts, of instance, in water content, temperature, microbial activity, and nutrient availability that occur within hours to days of a rainfall or irrigation event [3], [4]. Consequently, available soil databases systematically under-represent the temporal dynamics that affect ecosystem health and agricultural productivity.

In environmental monitoring and precision agriculture, high-frequency observations and predictive modelling are critical for informed decision-making. Recent advances in environmental sensing have produced a new generation of “3A” (available, accessible, and affordable) instrumentation allowing continuous, high-frequency monitoring of key soil state variables (e.g., moisture, temperature, pH), across scales from individual horizons to entire field profiles [5]. That in addition to earth observation platforms (e.g., Sentinal 1 (SAR), Landsat-8/9) offer spatially continuous data across scales from individual pedons to entire watersheds at diverse climate zones. Despite these advancements, current soil information systems lack the integrated computational frameworks that can combine multimodal data streams into dynamic, continuously updated representations of soil processes.

Three fundamental obstacles currently preventing the advancement of monitoring, modelling, and assessing the soil dynamical systems. First, disconnected models and observations, where traditional models such as HYDRUS [6] and SWAP [7] operate as isolated simulations calibrated from sparse field measurements rather than continuously updated from live sensor data. Second, broken interoperability, where global soil databases, such as SoilGrids [8], Soil Survey Geographic Database (SSURGO), and European Soil Data Centre (ESDAC), use incompatible property names, units, depth conventions, and classification systems, making cross-platform integration a labour-intensive manual task. Third, a functional inference-gap, wherein raw sensor signals (e.g., moisture, temperature, EC) fail to directly yield the high-order, agronomically meaningful variables, such as plant-available water, hydraulic conductivity, nutrient fluxes, and microbial respiration rates, that practitioners need for precise decision-making.

To addresses these obstacles, we present the Digital Pedon (DP) framework, a software system that represents the physical soil profile (pedon) as a dynamic digital twin. The DP framework is a computational object that ingests streams of multi-source sensor data and applies directed acyclic chain of built-in and user-defined solvers (Model-Zoo) to infer high-order (difficult to measure) soil variables in real-time. Interoperability is s achieved through an ontology layer aligned with FAO GLOSIS [9] and OGC SoilML standards [8], ensuring all properties carry machine-readable identifiers and standardized units. To ensure deployment flexibility, the DP framework is compatible with edge-computing systems (e.g., Raspberry Pi, Arduino, ESP32) and cloud geospatial platforms (e.g., Google Earth Engine, Microsoft Planetary Computer, Open Data Cube). By integrating real-time sensing, semantic data, and environmental modelling, the DP provides a scalable substrate for precision agriculture, digital soil mapping, and environmental health assessment, where the integration of diverse soil data with predictive modelling is increasingly critical.

## 2. Software description

### 2.1. Architecture

The DP package encompasses four modules: `pedon_api.py` (core DP framework), `ontology/vocab.py` (property vocabulary), `sources/soilgrids.py` (database connectors), and sensor/sensor_layer.py (BYOD sensor layer). Zero external runtime dependencies are required; optional `[yaml]` and `[llm]` extras add `PyYAML` and the Anthropic SDK respectively. Figure 1 illustrates the complete five-layer architecture and data flow. The architectural core separates concerns into three primary layers. The Structural Layer (SL), populated once via `build_pedon()` and never modified at runtime, stores horizon geometry (FAO/WRB [10]designation, `depth_top_cm`, `depth_bottom_cm`), 26 canonical soil properties, Van Genuchten hydraulic parameters [11], and threshold configuration. A Soil Type Registry (STR) provides five named templates (`loamy_topsoil`, `clay_subsoil`, `sandy_loam`, `peat`, `calcareous_loam`) via an inheritance-by-merge pattern. The Dynamic Layer (DL) manages live state through three optimised structures: a Key-Value Store (KVS) for *O*(1) current-state access, an append-only Time-Series Log (TSL) for temporal queries via `query_history()`, and a Threshold Event System (TES) that fires five agronomic alarms (saturation, wilting point, high/low temperature, high EC) through configurable observer-pattern callbacks.

**Figure 1.**
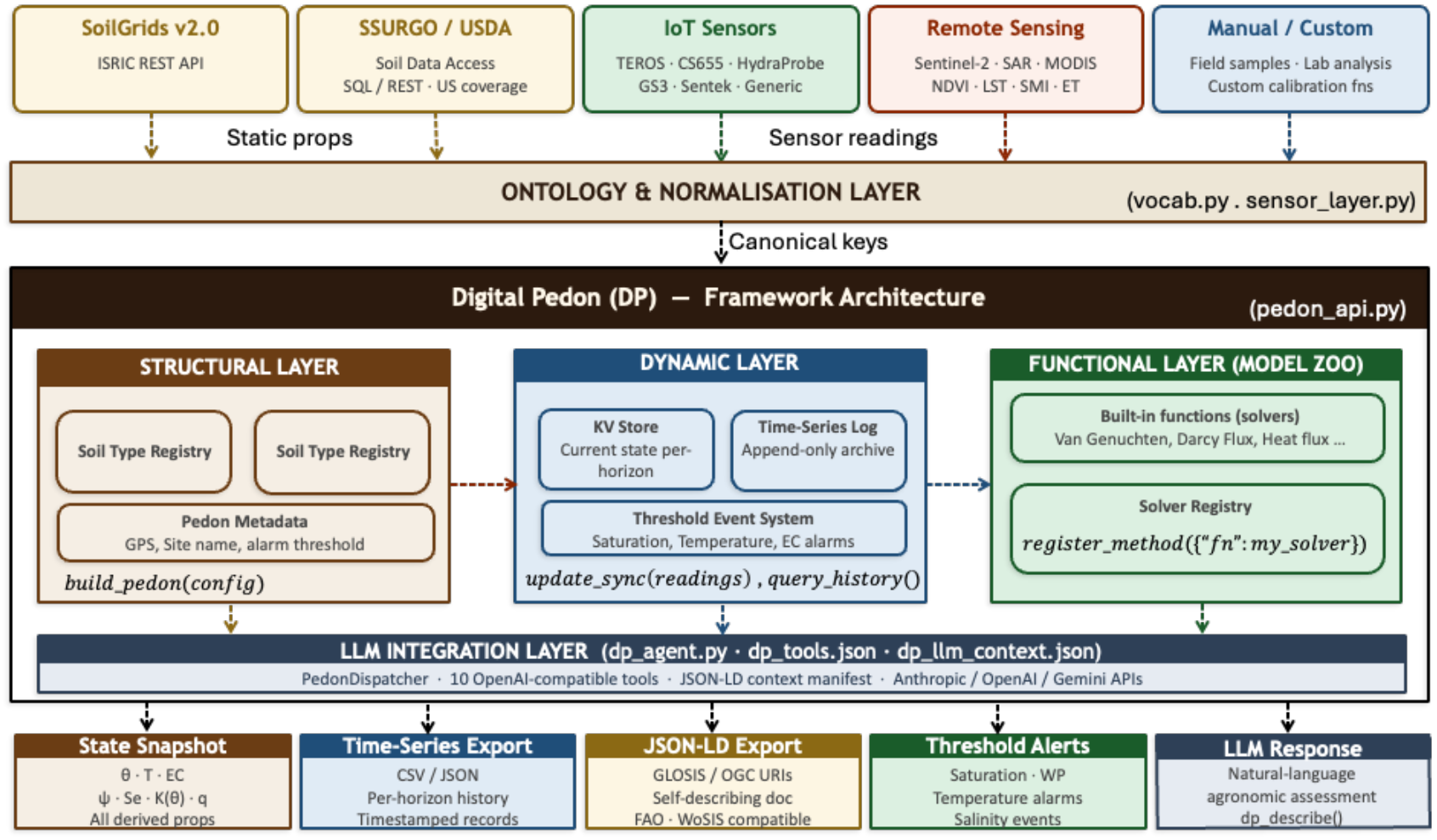
Digital Pedon (DP) framework architecture. Five input sources feed through the Ontology & Normalisation Layer into the three-layer core (Structural, Dynamic, Functional). The LLM Agentic Interface Layer exposes the framework to language models. Five output types are produced on every `update`() cycle.

The Functional Layer (FL) is a Model Zoo, where all registered solvers execute in sequence on every `update()` call, each receiving a params-dictionary and a state-dictionary and returning derived quantities that are merged into state, forming a Directed Acyclic Graph (DAG) of data dependencies. Two transversal layers complete the architecture, including the Ontology & Normalisation Layer (ONL) that resolves all incoming property names to GLOSIS URIs, and the LLM Agentic Interface Layer, which exposes the framework through 10 OpenAI-compatible tool schemas.

### 2.2. Software functionalities

The DP package provides seven core functionalities (F1-F7) designed to transition soil data from static records to dynamic, machine-interoperable digital twins:

> F1. Automated Initialization: The framework enables zero-configuration initialization through the `fetch_soil_profile(lat, lon`) method and GPS coordinates. It interfaces with the SoilGrids v2.0 REST API [8] and SSURGO to retrieve soil properties for six depth intervals. The DP automatically estimates Van Genuchten hydraulic parameters using Rawls-Brakensiek PTFs [10], allowing for a fully parameterized model without prior field work.
>
> F2. Real-time inference: At every `update()` call, the DP executes a solver chain, of built-in and user-defined models, as a suite of software sensors. The built-in solvers include Van Genuchten-Mualem [11] for moisture retention, Darcy-Buckingham [12] for vertical flux, a simplified De Vries model [13] for soil thermal properties, and a Q10 respiration [14] solver.
>
> F3. Extensible solver registry: The DP support an “open-world” approach to modelling. Users can register custom Python functions, to be added to the Model-Zoo, via `register_method()`. This enables complex inference chain, such as passing estimating matric potential from built-in solver into a custom root-uptake model, without modifying the core codebase. Built-in solvers can also be unregistered from the Model-Zoo and replaced with custom alternatives.
>
> F4. Bring-Your-Own-Device (BYOD) sensor manifest: To ingest multimodal data, the DP utilizes a sensor manifest that maps raw streams to the framework’s canonical vocabulary. The system includes 14-unit conversions (e.g., (% → cm^3^/cm^3^, °F → °C, μS/cm → dS/m, kPa → cm H_2_O, Sentinel-2 DN → reflectance). Remote sensing products (NDVI, LST, SAR soil moisture index) use the same manifest interface, unifying in-situ and satellite data ingestion.
>
> F5. Semantic Interoperability and JSON-LD export: The `vocan.py` module anchors 26 canonical soil properties to GLASIS/FAO [15] and OGC SoilML URIs [16]. The `canonical_key()` method resolves any vendor alias in *O*(1) and `normalise_dict()` rewrites all keys in a single pass. The `to_jsonld()` method exports state snapshots as self-describing documents, ensuring compliance with global standards like WoSIS [8], [17] and ESDC. The total vocabulary covers 41 properties (26 in `vocab.py` + 15 sensor-specific in `sensor_layer.py`).
>
> F6. Threshold event system: The DP features a configurable alarm system for critical agronomic states, including saturation, reaching wilting point, and salinity thresholds. Custom event handlers can be registered to automate hardware responses, such as triggering irrigation or drainage valves, based on physics-derived conditions rather than raw sensor values.
>
> F7. LLM agentic interface: To facilitate natural language interaction, `dp_agent.py` exposes the pedon’s state and solvers as OpenAI-compatible tool schemas. This allows Large Language Model (LLM) agents to query the profile (e.g., asking “Is the *Ap* horizon under water stress?”) and receive structured, physics-validated answers, without the user writing any API code.

## 1. Illustrative examples

Four experiments (E1–E4) demonstrate the capabilities of each framework layer. These examples are fully reproducible with “`pip install -e.`” using the included “`examples/`” scripts. The DP framework handles all remote API calls and parameterization internally (i.e., not external data files are required).

E1. GPS initialization: By invoking the `fetch_soil_profile(lat, lon)` method for four geographically different sites with contrasting soil texture classes (Table 1), the DP framework successfully generated fully parametrized digital pedons in a single function call. As shown in Table 1, the results demonstrate high physical consistency with the UNSODA hydraulic atlas [18]: the Pedotransfer function (PTF-)estimated saturated water content (*θ*_*s*_) increases monotonically from sand to clay, while saturated hydraulic conductivity (*K*_*s*_) decreases by over two orders of magnitude. The pore-size distribution index (*n*) also correctly scales inversely with clay content, validating the framework’s ability to bootstrap accurate physics-ready profiles using only a coordinate pair.

**Table 1.** Van Genuchten parameters, saturated water content (θ_s_), residual water contents (θ_r_), and air-entry pressure (α), from Rawls-Brakensiek Pedotransfer function PTFs for four GPS-initialized sites (Ap horizon).

| Site | Texture | $\theta_r$ | $\theta_s$ | $\alpha$ (cm<sup>-1</sup>) | $n$ | $K_s$ (cm/d) |
| --- | --- | --- | --- | --- | --- | --- |
| Ghent, Belgium | Loam | 0.034 | 0.480 | 0.020 | 1.41 | 15.0 |
| Niger, Sahel | Sand | 0.018 | 0.420 | 0.058 | 1.89 | 48.2 |
| Nile Delta, Egypt | Clay | 0.071 | 0.510 | 0.006 | 1.10 | 0.8 |
| Po Valley, Italy | Silty loam | 0.029 | 0.465 | 0.014 | 1.53 | 9.6 |

E2. Dynamic Solver Chain: A synthetic drying-wetting cycle, volumetric water content *θ* = 0.08 to 0.43 cm^3^/cm^3^, was ingested into a loamy topsoil pedon. All four solvers ran automatically on each `update_sync()` call. As shown in Figure 2, the matric potential *ψ* increases monotonically with *θ, K*(*θ*) follows the characteristic S-shape, and CO_2_ flux peaks near *θ*_*opt*_ *≈* 0.26 then declines as the Q10 moisture-scaling function suppresses aerobic respiration under waterlogging.

**Figure 2.**
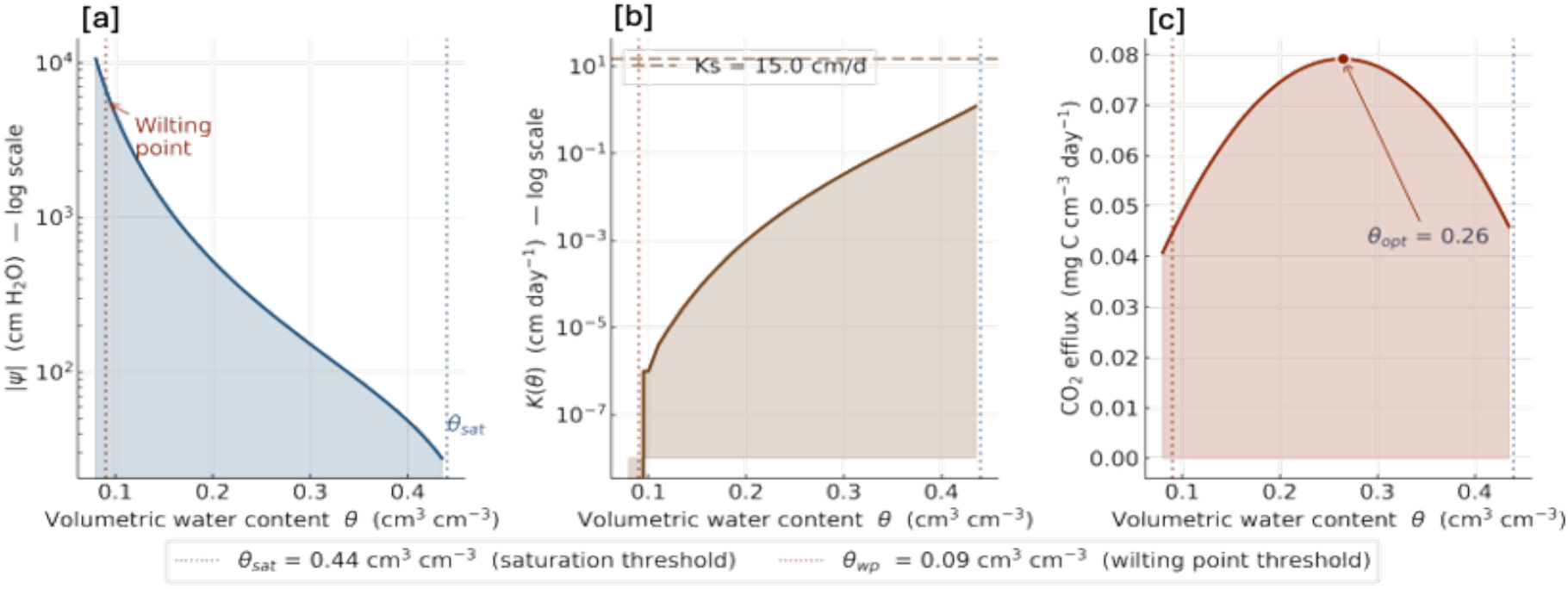
Solver chain output for a loamy topsoil across a drying–wetting cycle (θ = 0.08–0.43 cm^3^ cm^−3^, T = 15°C). All solvers ran automatically on each `update`() call: (a) matric potential |ψ| (Van Genuchten); (b) unsaturated hydraulic conductivity K(θ).

**Figure 3.**
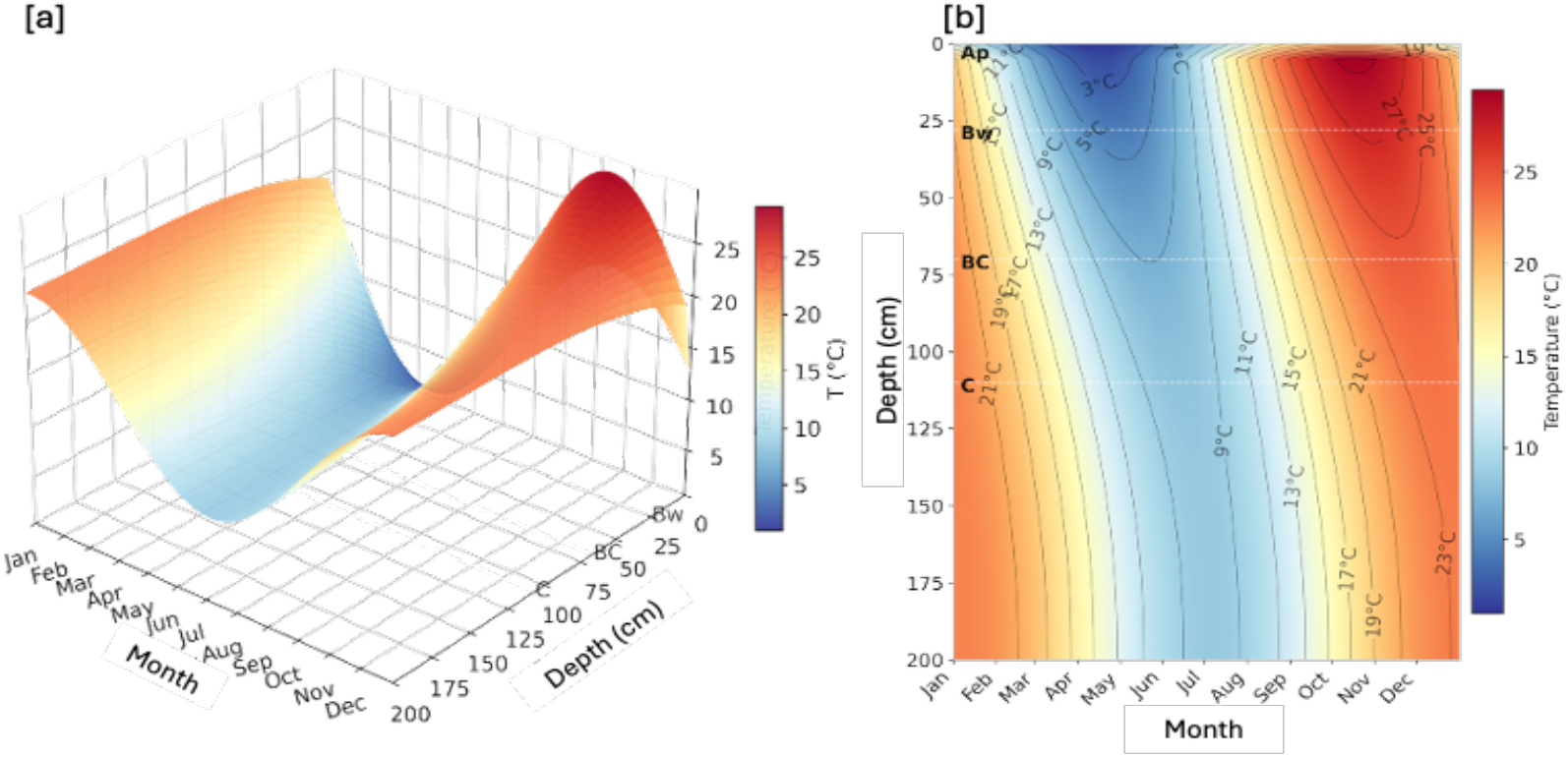
Seasonal soil heat conduction through a four-horizon loamy soil profile (Ghent, Belgium; 0–200 cm). Thermal diffusivity D_T_(z, t) is computed by the digital pedon’s De Vries built-in solver on every timestep (1 day), functioning as a user-defined solver utilizing built-in outputs. (a) A 3D temperature surface T(depth, time); (b) depth–time heatmap with isotherm contours (2°C interval). The dashed lines indicate the different horizon boundaries.

E3. Ontology compliance: To validate the framework’s semantic robustness, a comprehensive test suite (`pytest tests/test_ontology.py -v`) is implemented to evaluate the mapping accuracy of 28 property aliases across four major database conventions, namely, SSURGO, SoilGrids, ESDAC, and DONESOL. The experiment confirmed the reliability of the `normalise_dict()` round-trip using simulated SoilGrids payloads and verified the structural validity of the JSON-LD @context URIs. As shown in Table 2, all 28 tests passed successfully, with every generated URI correctly resolving to the w3id.org/glosis namespace.

**Table 2.** Representative alias resolution results (5 of 28 tests shown). Overall: 28/28 PASS.

| Source alias | Database | Canonical key | GLOSIS URI fragment | Result |
| --- | --- | --- | --- | --- |
| CLAY | SSURGO | clay_content | codelists#ClayContent | PASS |
| bdod | SoilGrids | bulk_density | codelists#BulkDensity | PASS |
| phh2o | SoilGrids | ph_h2o | codelists#SoilReactionpH | PASS |
| SOC_mean | ISRIC | organic_carbon | codelists#OrganicCarbon | PASS |
| VWC | Generic | volumetric_water_content | codelists#VolumetricWaterContent | PASS |

E4. User-defined solver extension: To demonstrate framework’s extensibility, a seasonal heat conduction model is implemented via `register_method()` without any modification in the core code. The solver implements the Fourier equation 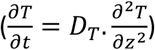 using a Crank-Nicolson [19] implicit scheme, with Δ*z* = 5cm and Δ*t* =1day, applied to a four-horizon loamy profile (Ap, Bw, BC, C; 0 – 200 cm) at Ghent, Belgium. The surface boundary was forced with an annual temperature cycle with mean *T* = 10.5°C and amplitude = 9.5°C. Unlike standalone simulations, the thermal diffusivity (*D*_*T*_) is not defined here as fixed parameter, instead, it is retrieved dynamically from the built-in De Vries solver on every timestep at each horizon node.

The user-defined solver thus used the output of a built-in solver within the same update cycle, demonstrating the compositionality of the DP framework’s directed acyclic solver chain. The validation against physically expected signatures confirmed the user-defined solver’s fidelity. The isotherm inflections visible at horizon boundaries in Figure 4 reflect the *D*_*T*_ contrast between the clay subsoil (Bw, 196–205 cm^2^ day^−1^) and the sandy BC horizon (152–162 cm^2^ day^−1^), confirming that the user-defined solver correctly inherits horizon-scale heterogeneity from the framework’s structural layer.

## 2. Impact

The novel framework of DP advances soil science by treating the pedon as a continuously up-to-date computational object rather than a static database record. Unlike conventional soil information systems (e.g., [8], [15]), the DP accommodates dynamic changes in soil state variables, enabling real-time investigation and monitoring of different soil properties such as hydraulic dynamics, inter-horizon heat transfer, and biologically mediated processes including microbial respiration and nutrient fluxes [20], [21], [22].The framework integrates multimodal data from in-situ sensor networks and satellite observations, allowing researchers to directly link remotely sensed indicators such as NDVI, land surface temperature, SAR-derived soil moisture, with subsurface soil processes. This integration improves digital soil mapping by enabling dynamic, sensor-driven updates of soil attributes instead of static simulations, enabling predictive monitoring and real-time risk simulation. This task is achieved through observational data streams that support solver-based inference methods. Unlike traditional models such as HYDRUS, which rely on fixed parameters and manual calibration, the DP framework utilizes solver chains within a continuously updated soil state representation to bridge data streams with process-based simulations. Furthermore, the DP substantially reduces operational overhead by automating tasks such initialising soil profiles from GPS coordinates and integrating sensor data streams, through the BYOD manifest layer, without extensive preprocessing. This allows scientists to focus on hypothesis testing and model development rather than data engineering. The framework’s compatibility with standard Python environments and its zero-dependency core enable straightforward deployment across infrastructures ranging from Raspberry Pi edge nodes to cloud geospatial platforms. Additionally, in fields such as precision agriculture, this translates directly into applications such as automated irrigation scheduling, real-time crop water stress assessment, and soil health monitoring at field scale. At broader scales, the DP’s scalable, ontology-compliant architecture supports soil carbon accounting, land degradation monitoring, and climate adaptation assessments, positioning it as a practical substrate for next-generation multi-platform digital agriculture systems.

An additional a unique capability of the DP is its LLM Agentic Interface Layer, which exposes the framework via OpenAI-compatible tool schemas and GLOSIS JSON-LD manifests, allowing models like Claude, GPT-4, and Gemini to query soil profiles (pedons) using natural language. This design transforms the DP into an AI-ready substrate for agronomic and ecological advisory systems. Such capability, which is currently not served by any existing soil modelling tool, allows practitioners to interrogate the pedon conversationally (e.g., “Is the Ap horizon under water stress?”) to receive structured, model-based insights without writing code.

## 3. Conclusions

This work introduces the Digital Pedon (DP), an open-source Python framework that models soil profiles as dynamic digital twins through three layers: a Structural Layer for immutable soil constants, a Dynamic Layer for real-time sensor data, and a Functional Layer for executing solvers. The DP resolves the three structural gaps in soil information systems, namely, the disconnected models and observations, cross-database interoperability, and the inference gap between raw sensor signals and agronomically meaningful variables. Four proof-of-concept experiments confirmed the framework’s correctness: GPS-xbased initialisation produced physically consistent Van Genuchten parameters across contrasting soil textures; the dynamic solver chain reproduced expected hydraulic, thermal, and respiratory responses; the ontology layer passed 28/28 alias mappings with full GLOSIS URI validity; and a user-defined Crank– Nicolson heat conduction solver confirmed solver compositionality and extensibility without core-code adaptation. The DP’s GLOSIS-compliant JSON-LD export supports direct WoSIS and ESDAC submission, while its LLM Agentic Interface Layer positions it as an AI-ready substrate for agronomic advisory systems. The framework is released under CC BY 4.0 at https://github.com/Pierianspring/digital-pedon. Future research will target scaling the framework from individual pedons to landscape-level digital twins, integrating novel data sources including soil spectroscopy, fused satellite multi/hyper-spectral imagery, and IoT sensor networks, thus presenting the surface and subsurface soils as dynamic factors in local ecosystems and improving real-time monitoring capabilities within agroecosystems.

## Declaration of generative AI use

During the preparation of this work, the authors used Claude (Anthropic) for two purposes: (1) grammar check and improving the clarity of selected manuscript passages; and (2) reviewing and debugging the Python source code and attached documentation. All scientific content, experimental design, physical interpretations, and conclusions were developed and verified entirely by the authors. All AI-assisted code was independently tested and validated against expected physical behaviour before inclusion. The authors take full responsibility for the integrity, accuracy, and reproducibility of both the manuscript and the released software.

## References

[1] S. D. Keesstra et al., ‘The significance of soils and soil science towards realization of the United Nations sustainable development goals,’ SOIL, vol. 2, no. 2, pp. 111–128, 2016, doi: 10.5194/soil-2-111-2016.

[2] B. Minasny et al., ‘Soil carbon 4 per mille,’ Geoderma, vol. 292, no. 4, pp. 59–86, Apr. 2017, doi: 10.1016/j.geoderma.2017.01.002.

[3] H. Vereecken et al., ‘Modeling Soil Processes: Review, Key Challenges, and New Perspectives,’ Vadose Zone Journal, vol. 15, no. 5, pp. 1–57, May 2016, doi: 10.2136/vzj2015.09.0131.

[4] A. B. McBratney, M. L. Mendonça Santos, and B. Minasny, ‘On digital soil mapping,’ Geoderma, vol. 117, no. 1–2, pp. 3–52, Nov. 2003, doi: 10.1016/S0016-7061(03)00223-4.

[5] R. A. Viscarra Rossel and J. Bouma, ‘Soil sensing: A new paradigm for agriculture,’ Agric. Syst., vol. 148, pp. 71–74, Oct. 2016, doi: 10.1016/j.agsy.2016.07.001.

[6] J. Šimůnek, M. T. Van Genuchten, M. Šejna, J. Šimůnek, and V. Zone, ‘Recent Developments and Applications of the HYDRUS Computer Software Packages,’ Vadose Zone Journal, vol. 15, no. 7, pp. 1–25, Jul. 2016, doi: 10.2136/vzj2016.04.0033.

[7] R. P. Bartholomeus et al., ‘SWAP 50 years: Advances in modelling soil-water-atmosphere-plant interactions,’ Agric. Water Manag., vol. 298, p. 108883, Jun. 2024, doi: 10.1017/wat.2023.4.

[8] L. Poggio et al., ‘SoilGrids 2.0: Producing soil information for the globe with quantified spatial uncertainty,’ SOIL, vol. 7, no. 1, pp. 217–240, Jun. 2021, doi: 10.5194/soil-7-217-2021.

[9] R. Palma et al., ‘GloSIS: The Global Soil Information System Web Ontology,’ Semantic Web: – Interoperability, Usability, Applicability, vol. 16, no. 5, May 2025, doi: 10.1177/22104968251363767.

[10] N. G. Patil and S. K. Singh, ‘Pedotransfer Functions for Estimating Soil Hydraulic Properties: A Review,’ Pedosphere, vol. 26, no. 4, pp. 417–430, Aug. 2016, doi: 10.1016/S1002-0160(15)60054-6.

[11] M. Th. van Genuchten, ‘A Closed-form Equation for Predicting the Hydraulic Conductivity of Unsaturated Soils,’ Soil Science Society of America Journal, vol. 44, no. 5, pp. 892–898, Sep. 1980, doi: 10.2136/sssaj1980.03615995004400050002x.

[12] G. Sposito, ‘General Criteria for the Validity of the Buckingham-Darcy Flow Law,’ Soil Science Society of America Journal, vol. 44, no. 6, pp. 1159–1168, Nov. 1980, doi: 10.2136/sssaj1980.03615995004400060006x.

[13] D. A. DeVries, ‘Thermal Properties of Soils,’ in Physics of Plant Environment, W. R. van Wijk, Ed., Amsterdam: North-Holland Publishing Co., 1963, pp. 210–235.

[14] H. Liu, F. Rezanezhad, Y. Zhao, H. He, P. Van Cappellen, and B. Lennartz, ‘The apparent temperature sensitivity (Q10) of peat soil respiration: A synthesis study,’ Geoderma, vol. 443, no. 7, p. 116844, Mar. 2024, doi: 10.1016/j.geoderma.2024.116844.

[15] R. Palma et al., ‘GloSIS: The Global Soil Information System Web Ontology,’ Semantic Web: – Interoperability, Usability, Applicability, vol. 16, no. 5, May 2025, doi: 10.1177/22104968251363767.

[16] D. G. Leibovici et al., ‘Geospatial standards: An example from agriculture,’ The Routledge Handbook of Geospatial Technologies and Society, pp. 60–75, Aug. 2023, doi: 10.4324/9780367855765-7.

[17] N. H. Batjes, E. Ribeiro, and A. Van Oostrum, ‘Standardised soil profile data to support global mapping and modelling (WoSIS snapshot 2019),’ Earth Syst. Sci. Data, vol. 12, no. 1, pp. 299–320, Feb. 2020, doi: 10.5194/essd-12-299-2020.

[18] A. Nemes, M. G. Schaap, F. J. Leij, and J. H. M. Wösten, ‘Description of the unsaturated soil hydraulic database UNSODA version 2.0,’ J. Hydrol. (Amst)., vol. 251, no. 3–4, pp. 151–162, Oct. 2001, doi: 10.1016/S0022-1694(01)00465-6.

[19] D. M. Pérez et al., ‘Temperature and water content estimation in soils of the semi-arid region of Brazil using finite difference and CFD,’ Eur. J. Soil Sci., vol. 75, no. 5, p. e13583, Sep. 2024, doi: 10.1111/ejss.13583.

[20] T. E. Ochsner, T. J. Sauer, and R. Horton, ‘Soil heat storage measurements in energy balance studies,’ Agron. J., vol. 99, no. 1, pp. 311–319, Jan. 2007, doi: 10.2134/agronj2005.0103S.

[21] B. Bond-Lamberty and A. Thomson, ‘Temperature-associated increases in the global soil respiration record,’ Nature 2010 464:7288, vol. 464, no. 7288, pp. 579–582, Mar. 2010, doi: 10.1038/nature08930.

[22] J. P. Schimel and J. Bennett, ‘NITROGEN MINERALIZATION: CHALLENGES OF A CHANGING PARADIGM,’ Ecology, vol. 85, no. 3, pp. 591–602, Mar. 2004, doi: 10.1890/03-8002.

